# Sequence-based pH-dependent prediction of protein solubility using CamSol

**DOI:** 10.1101/2022.05.09.491135

**Authors:** Marc Oeller, Ryan Kang, Pietro Sormanni, Michele Vendruscolo

**Affiliations:** Centre for Misfolding Diseases, Yusuf Hamied Department of Chemistry, University of Cambridge, Cambridge, UK

## Abstract

Solubility is a property of central importance for the use of proteins in research and in applications in biotechnology and medicine. Since experimental methods for measuring protein solubility are resource-intensive and time-consuming, computational methods have recently emerged to enable the rapid and inexpensive screening of large libraries of proteins, as it is routinely required in development pipelines. Here, we describe the extension of one of such methods, CamSol, to include in the predictions the effect of the pH of the solubility. We illustrate the accuracy of the pH-dependent predictions on a variety of antibodies and other proteins.

## Introduction

Solubility is a key property that underpins the developability potential of proteins in industrial pipelines^1–8^. Other such properties include expression yield, immunogenicity, chemical and conformational stability, propensity to self-associate, viscosity and polyspecificity^1–8^. Although proteins have evolved to be soluble to the level required to be functional in the cellular environment^4,9,10^, protein reagents for research, diagnostic and especially therapeutic purposes are commonly required to withstand the high concentrations necessary for storage and for certain administration routes such as subcutaneous delivery. This consideration implies that in most cases protein solubility must be optimised even beyond typical natural levels, and that specific formulation conditions must be identified to maximise solubility and stability of the product.

The solubility of proteins is defined in terms of the critical concentration at which soluble and insoluble phases are in equilibrium^2^. Solubility is therefore dependent on the formulation conditions, including in particular the storage temperature and the pH of the solution. As a consequence, formulation optimisation is a key step in drug development pipelines, as it is important to find the most suitable pH to ensure that the protein is sufficiently stable. Therefore, it is critical to optimise the solubility of proteins to maintain activity in storage and high concentration formulations.

Although several methods have been developed for the experimental measurement of protein solubility^2,5^, these are not readily amenable to high-throughput screenings, which are required to assess the large number of candidates typically available the early stages of industrial pipelines. For this reason, many computational prediction methods have been developed in recent years. PON-sol^11^, SOLpro^12^ and PROSO II^13^ use machine learning techniques such as support vector machines or random forest to predict solubility. Other methods, such as Solubis^14^ derive the solubility from aggregation prone regions calculated using the TANGO algorithm^15^. The use of molecular dynamics simulations to predict the solubility, such as in the case of the SAP method^16^, has also been reported.

In this work we generalise the CamSol method^17^, which was introduced to predict the solubility of protein variants, to predict the effects of varying the pH on protein solubility. Our approach encompasses three main features: (1) the calculation of partial charges using the Henderson-Hasselbalch equation, (2) the calculation of hydrophobicity values with pH-dependent logD values, and (3) the calculation of the context-dependent residue pKa values, either from the 3D structure when available^18,19^ or through a sequence-based prediction^20^ (**Figure 1**). By employing this method, we show that we can accurately predict the solubility behaviour at different pH values of proteins with varying sizes, including nanobodies, fulllength antibodies and intrinsically disordered proteins.

**Figure 1.**
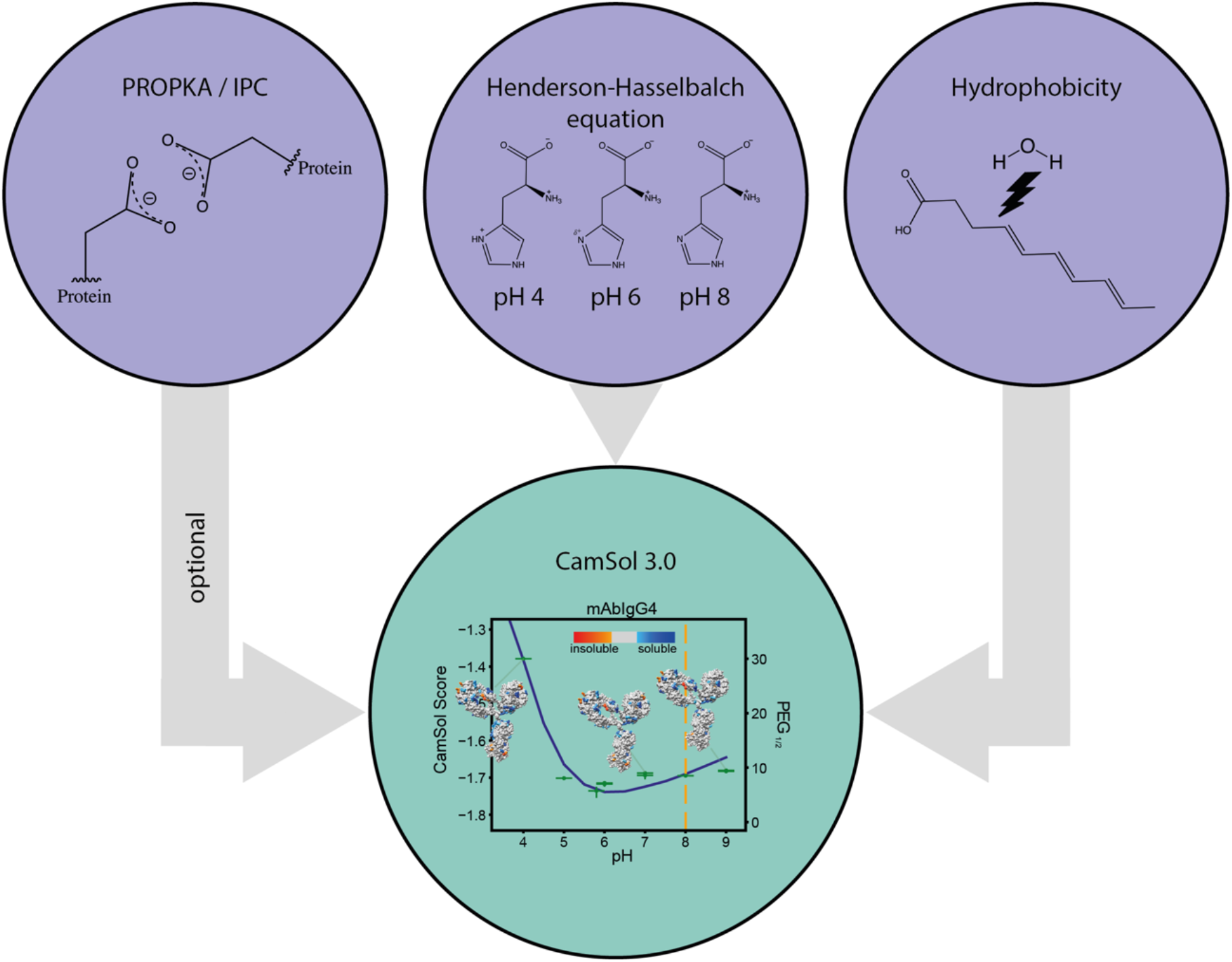
Schematic illustration of the sequence-based pH-dependent solubility predictions of CamSol. CamSol assesses partial charges using the Henderson-Hasselbalch equation. Hydrophobicity calculations are replaced by LogD calculations^21^. If a structure is supplied, amino acids pKa values are calculated using PROPKA, otherwise the IPC method is used. Experimental data (green markers in lower circle) were generated using a recently developed PEG Assay.

## Methods

### Theoretical methods

We provide an overview of the CamSol method and explain the changes introduced to take into account the effect of the pH on protein solubility. For a detailed explanation of the CamSol approach, the reader is referred to the original CamSol paper^17^. In CamSol, four physico-chemical properties, namely, charge, hydrophobicity, α-helical propensity and β-sheet propensity are combined to assign a score to each amino acid, yielding a solubility profile which is then smoothed and corrected for hydrophobic-hydrophilic patterns and gatekeeper effects. From this profile, an overall solubility score is calculated.

In the original CamSol method it was already possible to provide the value of the pH as input. The consequence of changing the input pH was to adopt specific side-chain charges depending on tabulated pKa values for the twenty standard amino acid. For example, all histidine residues would acquire a charge of +1 if the input pH was below 6.5. While rooted on general physicochemical principles, this description of the pH-dependence of the solubility can be substantially improved by more accurate calculations of the pKa values of the amino acids in their context. Therefore, in the current work, we updated the pKa values tabulated in CamSol by compiling all experimentally determined pKa values from http://compbio.clemson.edu/pkad (SI). We also introduced charges for the amide group at the N-terminus and the carboxylic acid at the C-terminus. The charge is calculated by using the Henderson-Hasselbalch equation, and partial charges are now allowed

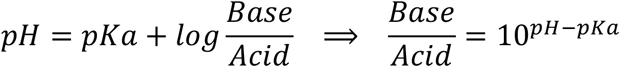

Using the ratio of charged to neutral species calculated with this equation, we replaced the logP values representing hydrophobicity by pH-dependent hydrophobicity values (logD). LogD combines the partition coefficient logP of neutral and ionised species

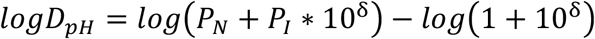

where δ is the difference between pKa and pH (pKa – pH for basic residues and pH – pKa for acidic residues). We used the pH-dependent LogD calculations by Zamora and colleagues^21^ for neutral and ionised LogP values for all standard amino acids. In the original CamSol method cysteine residues were assumed to be reduced. We changed the default to assume that all cysteine residues are in salt bridges and therefore cannot be charged. Free cysteine residues can be assigned by replacing it with the letter ‘B’ in the sequence.

To account for environmental effects that alter pKa values, we employed PROPKA, an opensource available pKa predictor^18,19^. When given a structure, PROPKA calculates pKa values for all ionisable residues based on their surrounding residues. If a structure or a suitable 3Dmodel is not available, as it is the case for disordered proteins and peptides, a sequencebased prediction is carried out instead. We employ IPC 2.0, which uses a mixture of support vector regression and deep learning models to predict pKa values based on the protein sequences^20^. Lastly, the pKa for histidine residues that are part of a terminal His-tag is lowered to 6 from 6.5 since the proximity of these histidine residues to each other causes a lowering of their pKa values to avoid electrostatic repulsion^22^.

Taken together, these changes enable the accurate prediction of the sequence-based pH-dependence of the solubility for a broad range of proteins.

## Experimental Methods

### Buffer

10 mM sodium phosphate dibasic heptahydrate (MP Biomedicals, 191441) and 10 mM citric acid monohydrate (Fisher Scientific, 5949-29-1) were combined. For each experiment the pH was adjusted using NaOH or HCl.

### Proteins

Bovine serum albumin (BSA, Sigma, A9418), human serum albumin (HSA, Sigma, A3782), and chicken egg white lysozyme (Sigma, L6876) were resuspended in buffer and then further purified and buffer-exchanged by carrying out size exclusion chromatography (SEC) (Cytiva, Superdex75 10/300 Increase for lysozyme and Superdex200 10/300 Increase for BSA and HSA). mAbIgG4 and its variant were provided by Novo Nordisk. DesAbO and α-synuclein were produced and purified in our group as described previously^23,24^. Each protein was freshly buffer-exchanged into the correct buffer (Cytiva, HiTrap Desalting column) before each assay.

### Solubility assay

The relative solubility of proteins was measured using a recently developed polyethylene glycol (PEG) solubility assay^25^. In brief, PEG 6000 is used as a crowding agent and titrated to final PEG concentrations of 0-33% and final protein concentration of 1 mg/mL, starting from PEG and protein stocks in the same buffer at the same pH, which typically needed to be re-adjusted after dissolving the PEG. After a 48 h incubation period, plates are centrifuged, the supernatant is transferred into a fresh plate and the soluble fraction is measured using a plate reader. A sigmoidal behaviour is seen for the precipitation of proteins, and the inflection point of the curve is used as a solubility proxy.

### Circular dichroism spectroscopy (CD)

Proteins were diluted to a concentration of 0.1 mg/mL. Three spectra were obtained using an Applied Photophysics Chirascan and a High Precision Cell made of quartz (Hellma Analytics, path length 1 mm) between 200 and 250 nm with a bandwidth of 1 nm, step size of 0.5 nm and scanning speed of 1 s/point.

### Spectrophotometry

Absorbance was measured with a plate reader (BMG Clariostar) and the spectrum from 220 to 700 nm was recorded at 25 °C.

## Results

The CamSol method calculates the solubility of proteins based on the physicochemical properties of their amino acid sequences^17^. Changes in pH mainly affect ionisable residues, as the pH determines the protonation state and therefore the electrostatic charges of these residues. To accurately assess the charge of each residue, we implemented the Henderson-Hasselbalch equation^26^ to determine the ratio between protonated and charged residues to estimate the partial charge of each amino acid.

An important component of these calculations is an accurate pKa value, which determines the pH range where the residue is charged. In this work, we employed three alternative strategies to obtain accurate pKa values. The user may choose which approach to use, depending on the information available on the protein under scrutiny. We revisited the tabulated pKa values used in the original version of CamSol for the 20 residue types and updated them with more accurate values.

In the first method, we retrieved pKa values from the computational biophysics and bioinformatics database (http://compbio.clemson.edu/pkad), which contains over 1500 experimentally measured residue pKa values from a wide range of proteins. Specifically, we employed the median pKa value observed in this database for each of the 20 amino acid types.

In the second method, sequence-based prediction of pKa values using IPC 2.0 were applied to all proteins with a specific focus on disordered proteins such as α-synuclein and peptides such as insulin. Using updated pKa values helps refine the solubility predictions and improves accuracy.

In the third method, we started from the observation that amino acid pKa values are affected by the structural environment and neighbouring residues. To take these effects into account we incorporated pKa predictions using the PROPKA method^18,19^. PROPKA is a widely used pKa predictor for proteins^27^, which requires the knowledge of the structure of the input protein.

Furthermore, we also ensured that the effect of changes in pH are accurately reflected in the way hydrophobicity affects the solubility predictions. Hydrophobicity so far has been expressed as the partition coefficient logP. To take charged species into account, we replaced these logP values with their corresponding logD values. Whereas logP values only consider the neutral state of a species, logD values take the neutral and charged states into account. We used data published by Zamora et al.^21^ to predict LogD values for all ionisable residues. Since logD values are also pKa-dependent, the calculated logD values are updated if a structure is available and PROPKA is used, and therefore the pKa values change. We normalised and fitted logD values to logP values previously used in CamSol to maintain consistency in the range of hydrophobicity values and to avoid the necessity to refit the scoring parameters of CamSol.

Since the availability of high-quality solubility data at varying pH values is limited, we set out to test our method on proteins that were either commercially available or already produced in our laboratory. A further criterion was that such proteins should not contain large cofactors (e.g. bound heme or large heteroatom groups) or metals as these can significantly alter their solubility and are not yet accounted for by the method. We measured the solubilities of α-synuclein, insulin, lysozyme, the single-domain antibody DesAbO^23^, HSA and BSA at pH values ranging from from 3 to 9 (**Figure S1**), as well as that of the full-length IgG4 monoclonal antibody HzATNP and one of its mutational variants^3^. To ensure that the solubility measurements are not affected by conformational changes of the proteins at certain pH values, we measured the CD spectra of every sample after carrying out the solubility assay (**Figure S2**).

We calculated the correlation between measured values and their predicted counterparts, testing also the different ways of calculating residue pKas implemented in the method (**Figure 2**). The results with IPC and without any pKa corrections are shown in **Figures S3 and S4**, respectively. Our results show that CamSol not only captures the overall trend in solubility upon pH changes well, but the predicted values are also highly correlated with the experimental values with Pearson’s coefficients of correlation always being greater than 0.67 (**Table S1**). **Figure 2a-d** depict predictions including PROPKA for folded, globular proteins, whereas **Figure 2e,f** (blue box) show the IDP α-synuclein and the peptide insulin for which IPC was applied. With the proteins tested experimentally, we tried to cover a broad range (4.5 – 11) of theoretical pI values (**Figure 2**, yellow vertical line), as the pI is usually a good indicator for the pH value corresponding to the minimum in solubility.

**Figure 2.**
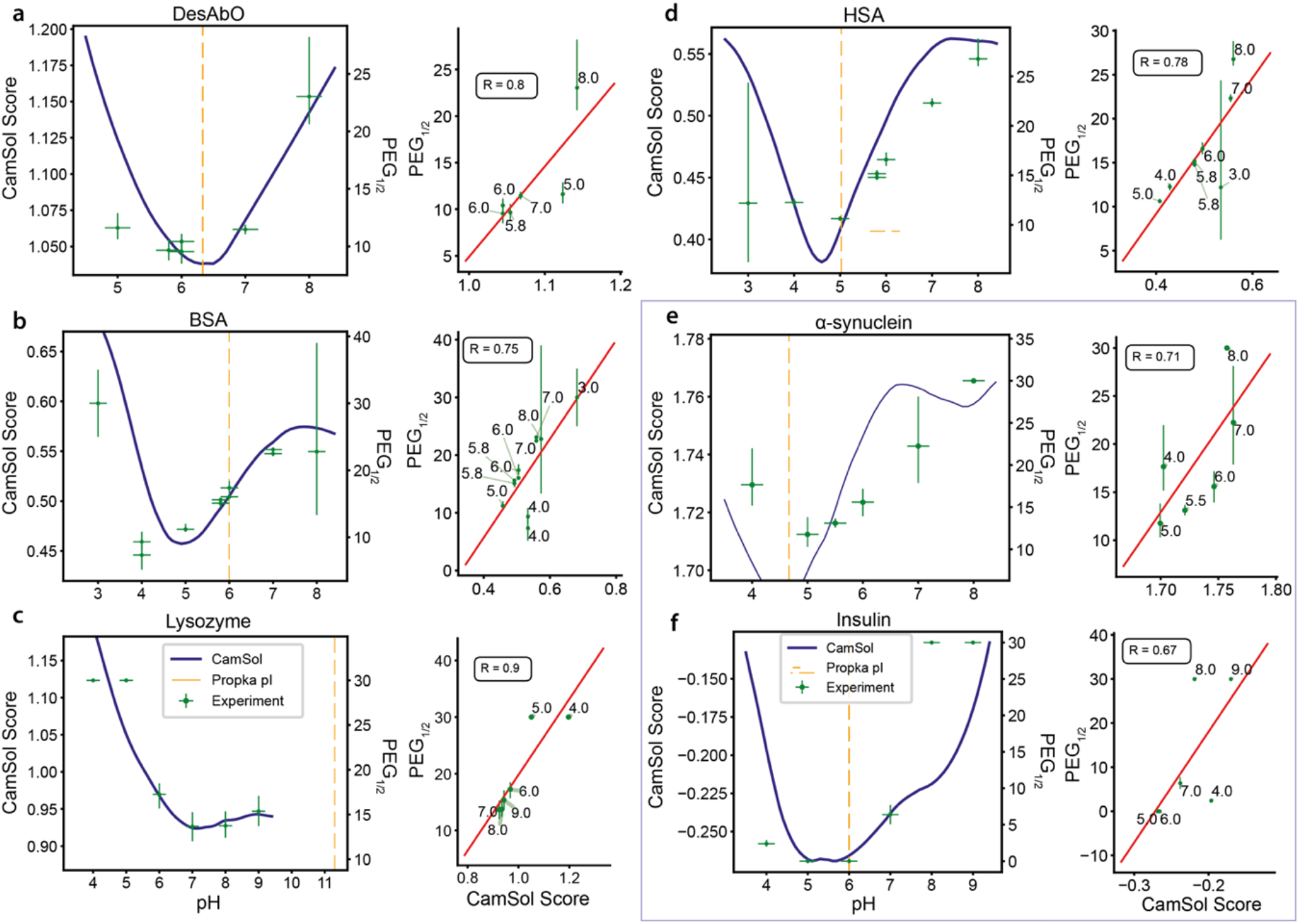
CamSol predicts solubility values that are highly correlated with experimental solubility values. Plots on the left-hand side in each column visualise how experimental and predicted values change over a range of pH values. The left axis and blue line report the predicted CamSol solubility score, the right axis and green markers the measured midpoints of PEG-precipitation, all as a function of pH (x-axis). The vertical yellow line is the theoretical isoelectric point. Plots on the right-hand side shows the correlation between the predicted and measured relative solubility values. CamSol calculations were carried out using pKa values calculated by PROPKA for **(a)** DesAbO (nanobody), **(b)** bovine serum albumin (BSA), **(c)** hen egg white lysozyme, and **(d)** human serum albumin (HSA), while for **(e)** α-synuclein and **(f)** insulin pKa values were calculated with IPC (framed in blue box). R is the Pearson’s coefficient of correlation.

BSA and HSA (**Figure 2b,d**) show good agreement between experimental values and predicted results with a Pearson’s coefficient of correlation of 0.75 and 0.78, respectively. These correlations improve slightly with the use of IPC to 0.92 and 0.88 and are still relatively good even if the pKa values are not corrected (0.63 and 0.78). The nanobody shows an even better agreement with a coefficient of correlation of 0.8. However, the case of the nanobody indicates the limits of the IPC approach, as the coefficient of correlation drops to 0.47 (0.39 without any correction). Lysozyme shows almost a perfect correlation, with a correlation coefficient of 0.9. For α-synuclein and insulin only the IPC method was applied since IDPs and peptides do not form stable structures that can be used for PROPKA. The correlations are slightly lower, with a coefficient of correlation of 0.71 and 0.67, respectively (0.72 and −0.67 without any correction).

We then tested the method on a IgG4 antibody (mAbIgG4). The coefficient of correlation is 0.96 (**Figure 3a**), 0.88 for IPC (**Figure S5**) and 0.85 for no correction (**Figure S6**). We visualised the change in solubility by projecting the solubility profile onto the structure of the antibody (blue: highly soluble region, red: highly insoluble region). It highlights how some regions become less soluble at higher pH values and vice versa. Following these results, we tested a variant of the antibody with a slightly lower pI to see whether CamSol was capable of capturing even small changes in pH-dependent solubility induced by minor mutations. CamSol can predict well the solubility of this variant with a coefficient of correlation of 0.97 (0.86 and 0.82 for IPC and no pKa correction) (**Figure 3b**).

**Figure 3:**
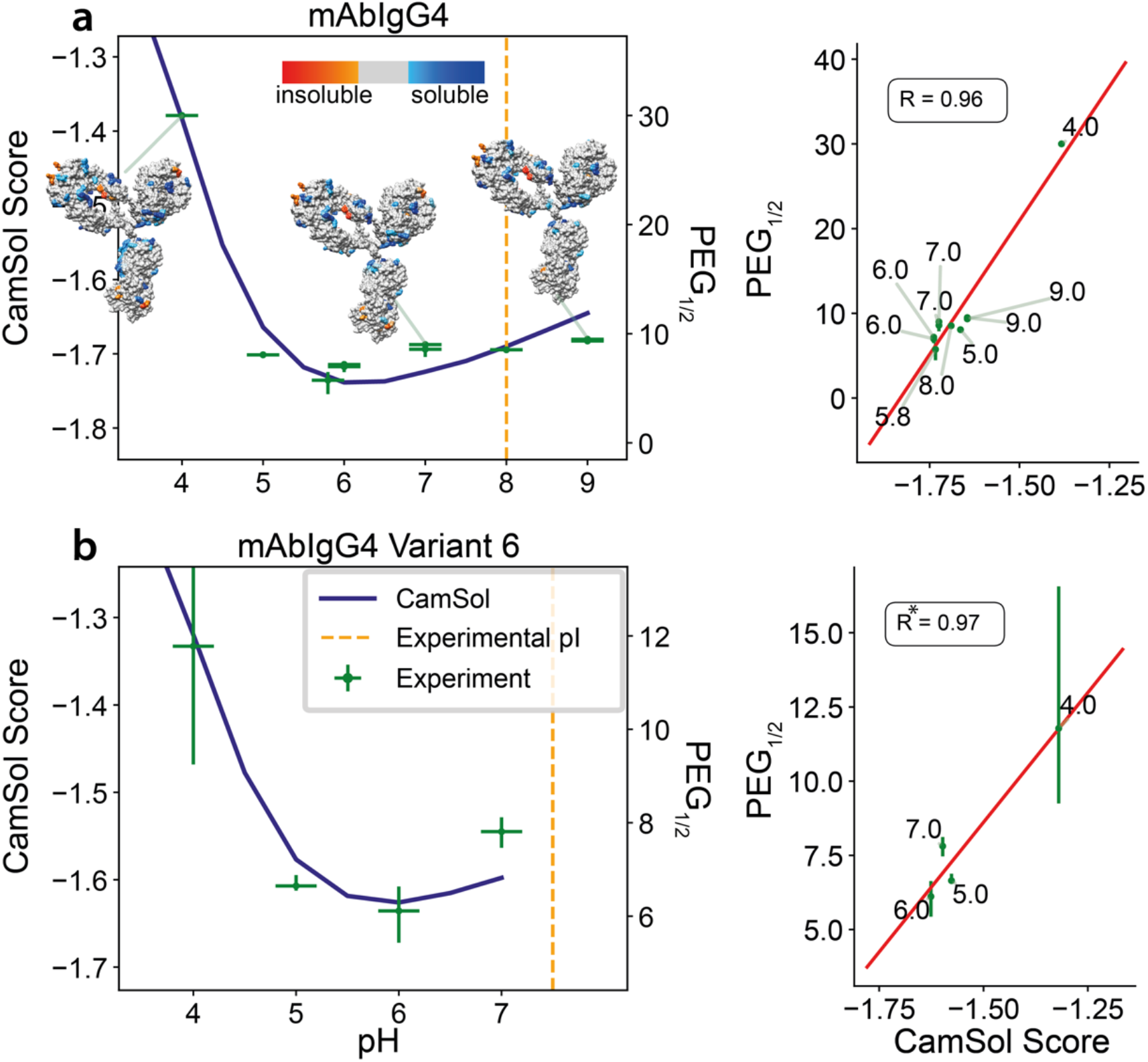
Change in solubility at varying pH for a monoclonal IgG4 antibody. As in Figure 2, plots on the left visualise how experimental and predicted values change over a range of pH values, and those on the right show the correlation between the predicted and measured relative solubility values. In **(a) t**he modelled structures of the IgG4 antibody are colour-coded according to the CamSol structurally corrected profile for pH 4, 7 and 9 (blue: highly soluble region, red: highly insoluble region, see colour-bar). **(b)** Same plot for a mutational variant of mAbIgG4 harbouring two mutations that lower the pI (S70E V99D on heavy and light chain respectively). R is the Pearson’s coefficient of correlation. R* correlation is given for consistency but is only an estimate since only four points are available. Green crosses: experimental values; blue lines: predicted values; yellow dotted line: calculated pI; green dotted line: experimental pI.

When we analysed DesAbO, we realised that the charge effect of the C-terminal His-tag (i.e. the sequence of seven histidine residues used for purification) could be estimated more accurately. The His-tag was not part of the structure and the pKa values were therefore not adjusted by using PROPKA. Nevertheless, it is clear from a physico-chemical standpoint that the pKa values must change if so many ionizable residues are near each other. By assuming the pKa value shifts from 6.5 to a lower value^22^, we carried out the calculations at pKa 6.

## Discussion and Conclusions

Predicting the solubility of proteins as a function of the pH is of great importance in industrial pipelines as it can reduce the number of cost and labour-intensive in vitro assays to determine optimal formulations^1,3–5^. With the new version of CamSol that we reported here we took a step into the that direction.

We have shown in this work that CamSol can now reliably predict the solubility of globular proteins with a wide range of pI values (from 4.5 for α-synuclein up to 11 for lysozyme). We showed that CamSol is also capable of capturing the changes induced by small mutations that shift the overall pI value of the protein. Using the new CamSol method we also visualised the changes of solubility on the surface of an IgG4 antibody, and demonstrated how certain areas of a protein can move from highly soluble to neutral and neutral areas to highly insoluble over a short range of pH values.

The pI value is usually used as an approximation of the pH value at which a protein is least soluble as this is the point at which the overall charge on a protein is neutral. Although in many cases this assumption is reasonably accurate, in general the solubility of folded proteins does not only depend on the overall charge. The distribution of these charges is crucial, as patches of opposing charges can act as starting points for aggregation due to strong attractive intermolecular forces. Although the changes in hydrophobicity are related to the changes in charge upon pH variations, these are not perfectly correlated and hence can move the point of least solubility away from the pI value as well. CamSol tries to capture all these effects to give a more accurate estimate of the point of lowest solubility. Moreover, we also predicted the behaviour around the pI value that is more informative than just the point of least solubility.

We have provided the user with three different options to predict the solubility: (1) Using tabulated pKa values, (2) Using PROPKA if a structure is available and the protein of interest is stably folded, to estimate the pKa values by assessing structural effects and improve the reliability of the predictions, and (3) using IPC, if no structure is available, to correct pKa values for neighbouring effects.

We acknowledge that the new version of CamSol is still limited for proteins that contain large co-factors such as heme, as we expect that these can alter the pH-dependent solubility significantly and CamSol cannot currently account for these aspects.

In conclusion, we have presented an extension of the CamSol method for the sequence-based prediction of the solubility of proteins at varying pH values.

## Acknowledgements

We thank Prof. Zamora for help and additional data provided to calculate LogD values. We also would like to thank Prof. Kozlowski for help implementing IPC 2.0. We are grateful to collaborators at Novo Nordisk (Dr Nikolai Lorenzen, Dr Marie Pedersen and Dr Daniele Granata) for providing the purified antibody HzATNP and its variant. M.O. is a PhD student funded by AstraZeneca. P.S. is a Royal Society University Research Fellow (URF\R1\201461).

## Author contribution

M.O. performed experiments and carried out data analysis. M.O. and R.K. purified alpha synuclein and measured its solubility. M.O. and P.S. wrote the software. M.O., P.S. and M.V. wrote the manuscript. M.O., P.S. and M.V. conceived and P.S. and M.V. supervised the project.

## References

1. Norman, R. A. et al. Computational approaches to therapeutic antibody design: established methods and emerging trends. Brief. Bioinform. 00, 1–19 (2019).

2. Wolf Pérez, A. M., Lorenzen, N., Vendruscolo, M. & Sormanni, P. Assessment of Therapeutic Antibody Developability by Combinations of In Vitro and In Silico Methods. in Methods in Molecular Biology 2313, 57–113 (2022).

3. Wolf Pérez, A. M. et al. In vitro and in silico assessment of the developability of a designed monoclonal antibody library. MAbs 11, 388–400 (2019).

4. Sormanni, P. & Vendruscolo, M. Protein Solubility Predictions Using the CamSol Method in the Study of Protein Homeostasis. Cold Spring Harb. Perspect. Biol. 1–12 (2019). doi:10.1101/cshperspect.a033845

5. Jain, T. et al. Biophysical properties of the clinical-stage antibody landscape. Proc. Natl. Acad. Sci. U. S. A. 114, 944–949 (2017).

6. Raybould, M. I. J. et al. Five computational developability guidelines for therapeutic antibody profiling. Proc. Natl. Acad. Sci. 116, 4025–4030 (2019).

7. Jarasch, A. et al. Developability assessment during the selection of novel therapeutic antibodies. J. Pharm. Sci. 104, 1885–1898 (2015).

8. Lauer, T. M. et al. Developability index: A rapid in silico tool for the screening of antibody aggregation propensity. J. Pharm. Sci. 101, 102–115 (2012).

9. Vecchi, G. et al. Proteome-wide observation of the phenomenon of life on the edge of solubility. Proc. Natl. Acad. Sci. 117, 1015–1020 (2020).

10. Tartaglia, G. G., Pechmann, S., Dobson, C. M. & Vendruscolo, M. Life on the edge: a link between gene expression levels and aggregation rates of human proteins. Trends Biochem. Sci. 32, 204–206 (2007).

11. Yang, Y., Niroula, A., Shen, B. & Vihinen, M. PON-Sol: Prediction of effects of amino acid substitutions on protein solubility. Bioinformatics 32, 2032–2034 (2016).

12. Magnan, C. N., Randall, A. & Baldi, P. SOLpro: Accurate sequence-based prediction of protein solubility. Bioinformatics 25, 2200–2207 (2009).

13. Smialowski, P., Doose, G., Torkler, P., Kaufmann, S. & Frishman, D. PROSO II - A new method for protein solubility prediction. FEBS J. 279, 2192–2200 (2012).

14. Ganesan, A. et al. Structural hot spots for the solubility of globular proteins. Nat. Commun. 7, 1–15 (2016).

15. Fernandez-Escamilla, A. M., Rousseau, F., Schymkowitz, J. & Serrano, L. Prediction of sequence-dependent and mutational effects on the aggregation of peptides and proteins. Nat. Biotechnol. 22, 1302–1306 (2004).

16. Chennamsetty, N., Voynov, V., Kayser, V., Helk, B. & Trout, B. L. Design of therapeutic proteins with enhanced stability. Proc. Natl. Acad. Sci. 106, 11937–11942 (2009).

17. Sormanni, P., Aprile, F. A. & Vendruscolo, M. The CamSol method of rational design of protein mutants with enhanced solubility. J. Mol. Biol. 427, 478–490 (2015).

18. Søndergaard, C. R., Olsson, M. H. M., Rostkowski, M. & Jensen, J. H. Improved treatment of ligands and coupling effects in empirical calculation and rationalization of p Kavalues. J. Chem. Theory Comput. 7, 2284–2295 (2011).

19. Olsson, M. H. M., Søndergaard, C. R., Rostkowski, M. & Jensen, J. H. PROPKA3: Consistent Treatment of Internal and Surface Residues in Empirical pka Predictions. J. Chem. Theory Comput. 7, 525–537 (2011).

20. Kozlowski, L. P. IPC 2.0: Prediction of isoelectric point and pKa dissociation constants. Nucleic Acids Res. 49, W285–W292 (2021).

21. Zamora, W. J., Campanera, J. M. & Luque, F. J. Development of a Structure-Based, pH-Dependent Lipophilicity Scale of Amino Acids from Continuum Solvation Calculations. J. Phys. Chem. Lett. 10, 883–889 (2019).

22. Qiagen. The QIA expressionist. Qiagen GmbH, Düsseldorf, Germany (2003).

23. Aprile, F. A. et al. Rational design of a conformation-specific antibody for the quantification of Aβ oligomers. Proc. Natl. Acad. Sci. 201919464 (2020). doi:10.1073/pnas.1919464117

24. Staats, R. et al. Screening of small molecules using the inhibition of oligomer formation in α-synuclein aggregation as a selection parameter. Commun. Chem. 3, 1–9 (2020).

25. Oeller, M., Sormanni, P. & Vendruscolo, M. An open-source automated PEG precipitation assay to measure the relative solubility of proteins with low material requirement. Sci. Rep. 11, 1–10 (2021).

26. Po, H. N. & Snozan, N. M. The Henderson-Hasselbalch Equation: Its History and Limitations. J. Chem. Educ. 78, 1499–1503 (2001).

27. Davies, M. N., Toseland, C. P., Moss, D. S. & Flower, D. R. Benchmarking pKa prediction. BMC Biochem. 7, 1–12 (2006).

