## Supplementary_Figures for "Sequence-based pH-dependent prediction of protein solubility using CamSol"

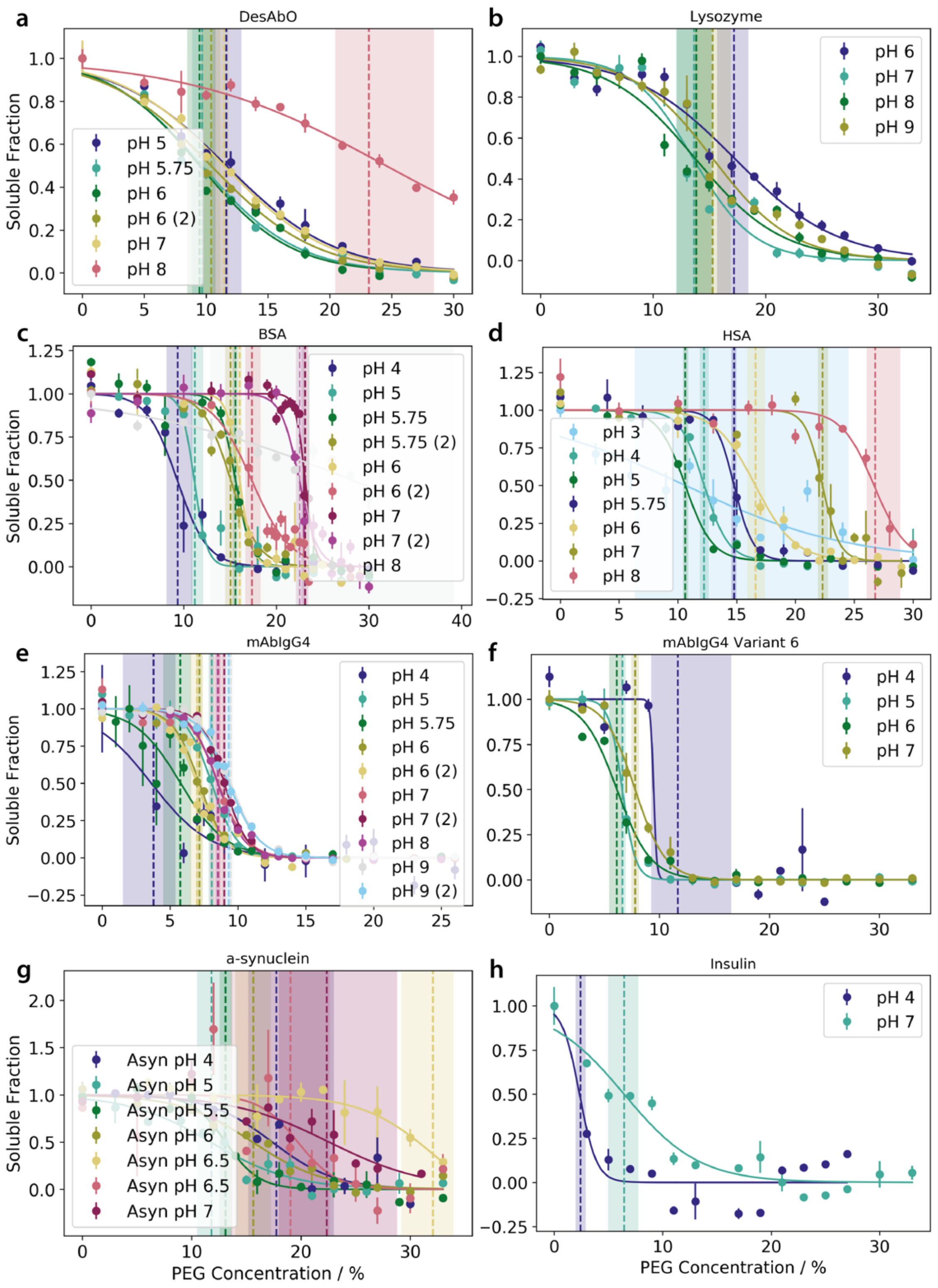


**Figure S1: Determination of the solubility of the proteins analysed in this work.** The solubility was measured using a recently developed PEG solubility assay. Measurements at pH values at which proteins were completely insoluble or completely soluble are not depicted for clarity.

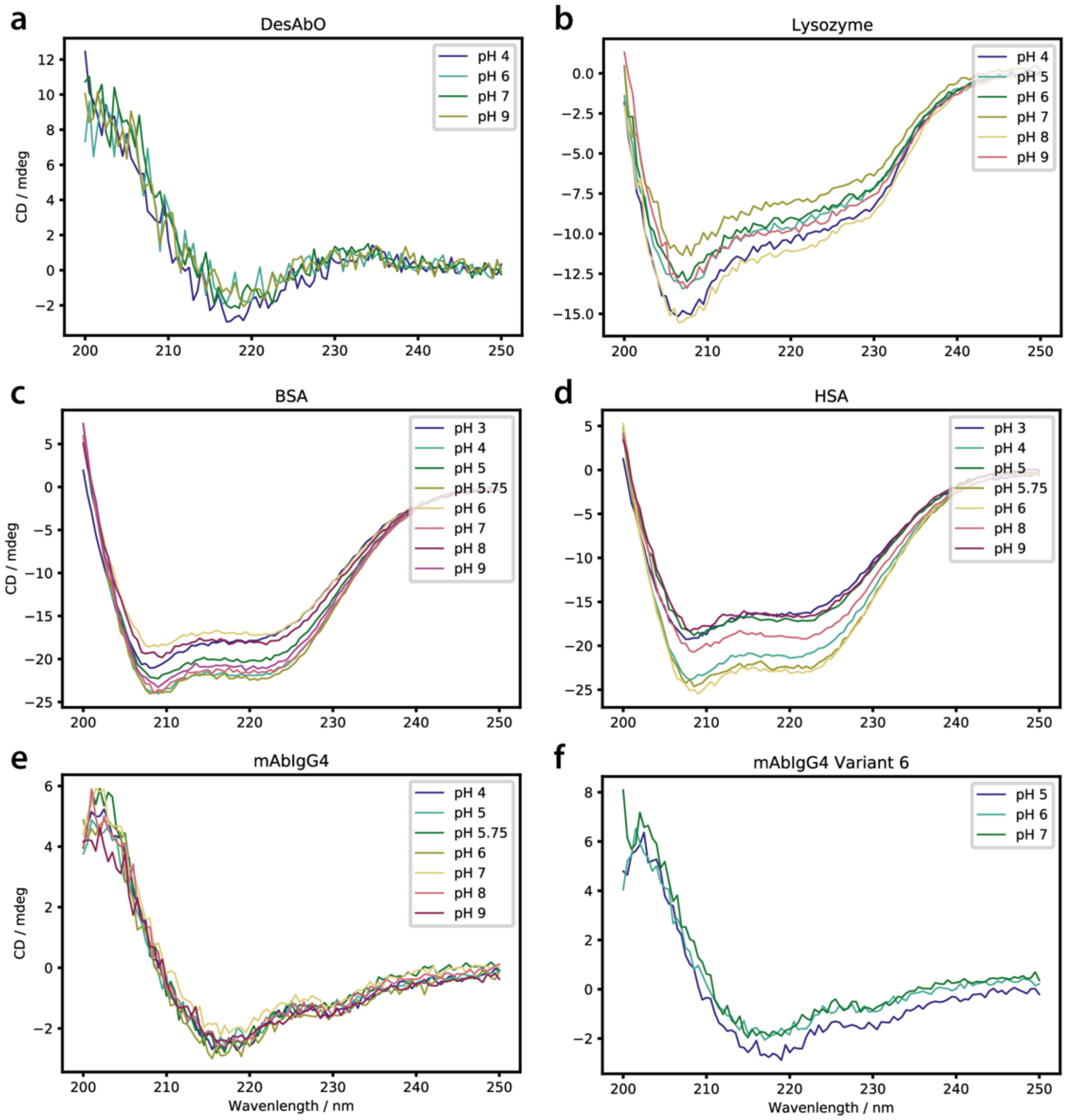


**Figure S2: CD spectra of all globular proteins used in this project.** CD spectra show that protein conformations are preserved across the pH ranges tested.

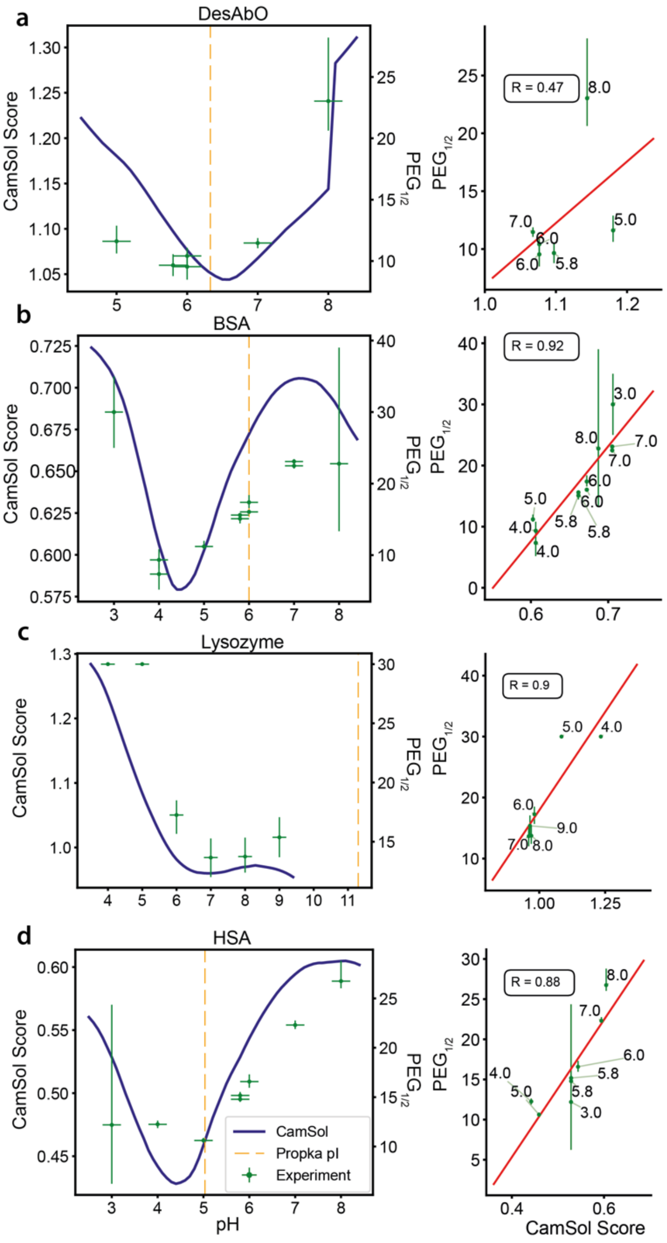


**Figure S3: Correlation of predicted and measured protein solubility with IPC used to correct pKa values.** Correlation for lysozyme is the same and for BSA and HSA is slightly higher than for case where PROPKA is used, however, the prediction of DesAbO is lower and below an acceptable threshold.

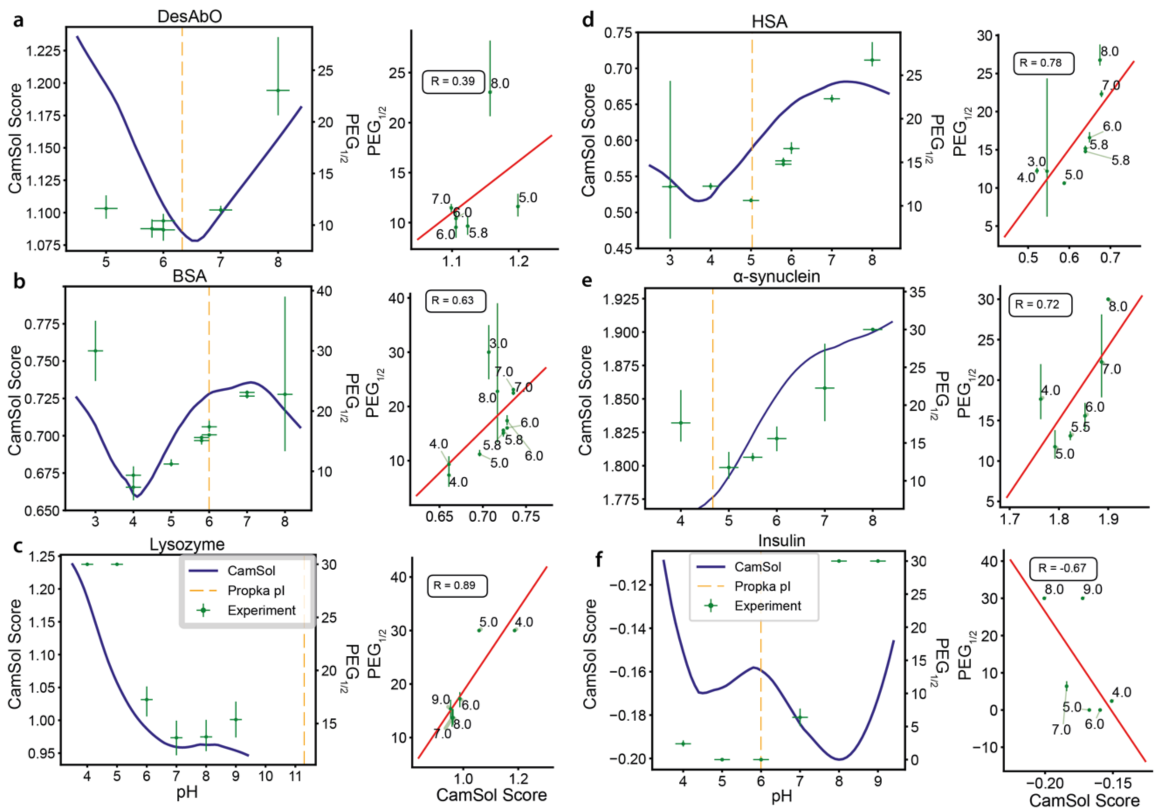


**Figure S4: Correlation between predicted and measured protein solubility without pKa corrections.** The correlation for lysozyme and HSA is the same and for BSA and α-synuclein is slightly lower than for case where pKa corrections is used. The predictions for insulin are not accurate, highlighting the importance of the pKa corrections.

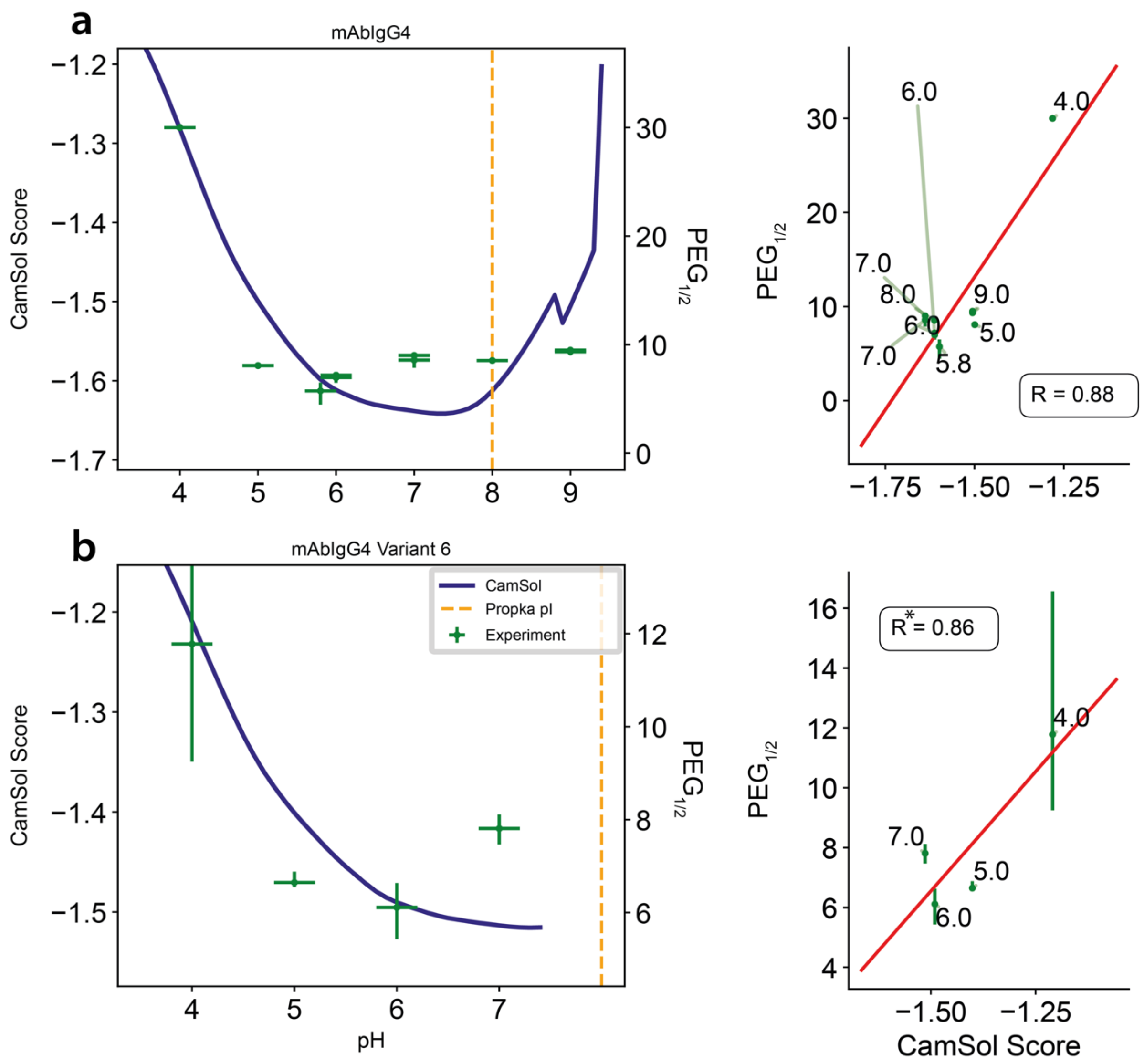


**Figure S5: Correlation of predicted and measured solubility of mAbIgG4 and its variant with IPC as the pKa correction method.** Correlation for the antibody and its variant is slightly smaller than for the PROPKA corrected case.

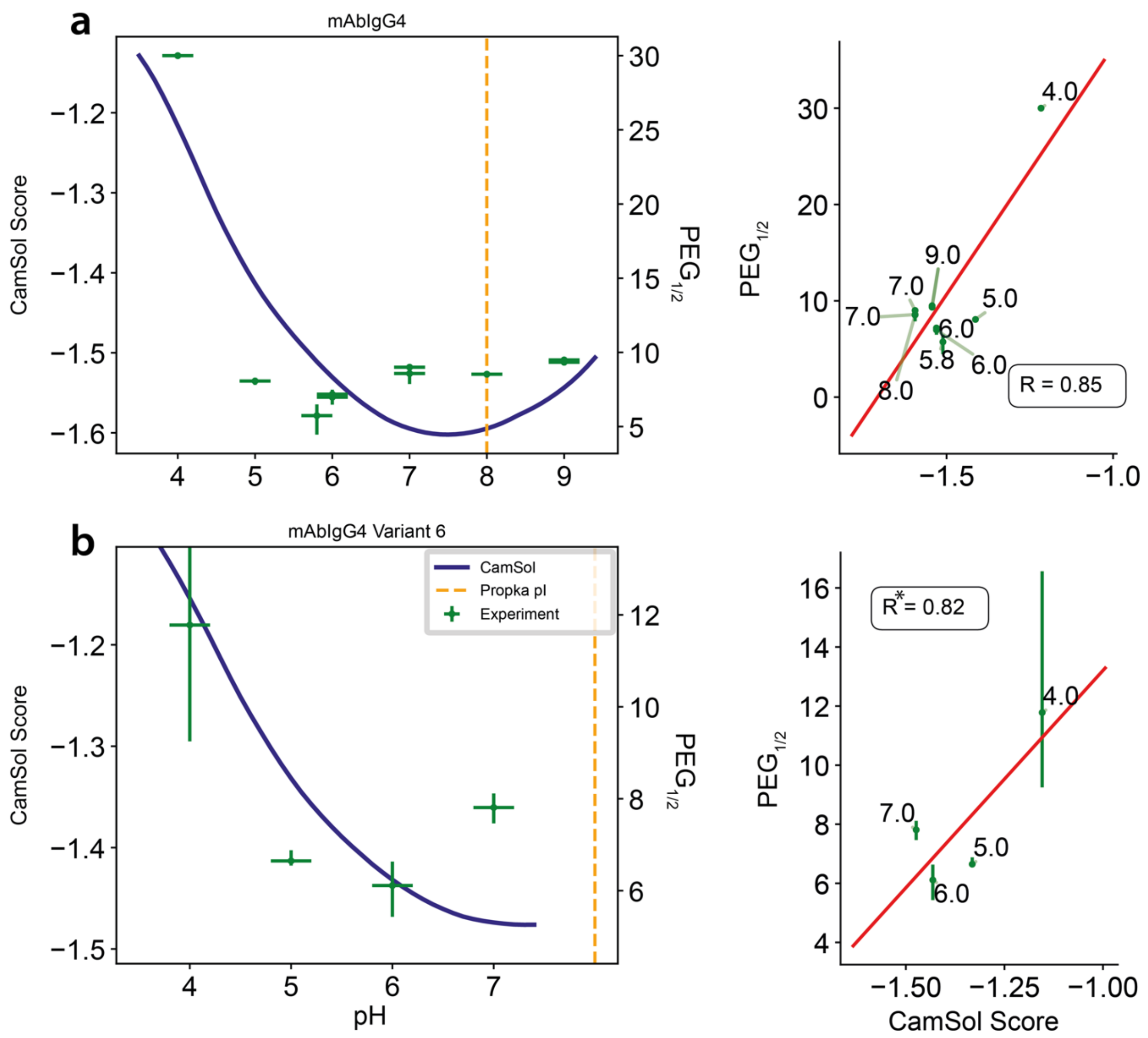


**Figure S6: Correlation of predicted and measured solubility of mAbIgG4 and its variant without pKa corrections.** Correlation for the antibody and its variant is slightly smaller than for the pKa corrected methods.
